# Comparative Nephro-Hepatoprotective Effects of Herbal Teas from *Hibiscus sabdariffa, Moringa oleifera, Zingiber officinale, and Azadirachta indica* in Alloxan-Induced Diabetic Male Wistar Rats: A Preclinical Study

**DOI:** 10.1101/2024.06.17.599331

**Authors:** Emieseimokumo Numonde, Isaac Sokoato Momoh, Victor Duniya Sheneni, Ebunoluwa Ajibike Okosesi, Micheal Omeyiza Ibrahim

## Abstract

Diabetes, a silent killer, ravages the kidney and liver, leaving a trail of destruction in its wake. Studies have suggested a linkconnectingdysfunctional liver and renalactivity along side glucotoxicretinopathy in patients of type 2 diabetes mellitus. Against this background, our preclinical study unveils the remarkable nephro-hepatoprotective effects of herbal teas from *Hibiscus sabdariffa, Moringa oleifera, Zingiber officinale, and Azadirachta indica*in diabetic male Wistar rats induced with alloxan.Diabetes of artificial source was administered through a one-time interperitonealadministeration of about seven categorieshaving 6 rats in the individual group.control (glycemic), diabetic groupthat is made up of alloxan treatment + Zobo (100+400 mg/kg)., alloxan + moringa (100 +200 mg/kg), alloxan + ginger (100+500 mg/kg), alloxan + Dogoyaro (100+250 mg/kg), and lastly, the alloxan +glibenclamide(100+5mg/kg) were orally given for 28 days.Reduction of tissue weight was observed upon administration of alloxan which was ameliorated upon treatment with the selected herbal teas. Also, elevated levels of liver and kidney biomakersinduced by alloxan were reversed upon administration of the Herbal teas. Furthermore, the Herbal teas decreased fasting blood glucose, which was initially significantly increased byalloxan (p<0.05). Consequently, given to their antihyperglycemic and nephro-hepatoprotective prowess, these selected herbal teas offer a beacon of hope for the millions afflicted not just with diabetes but also diabetes-related kidney and liver diseases.

## 1. Introduction

Diabetes is a chronic condition characterized by uneven metabolism of carbohydrate-containing diets that develop when the islet of Langerhans cells in the pancreas get disrupted by auto-immune mechanism and fails to produce insulin or when the cells of hepatic and extrahepatic tissues fail to take up glucose despite insulin availability. High blood sugar, increase in weight, incessant urination, dehydration, and constant hunger are all the symptoms. Diabetes is often classified based on clinical manifestation which are Type 1 diabetes which is common among young individuals, type 2 diabetes, and gestational diabetes.

Type 2 diabetes has the largest spectrum of occurrence and 90 percent of diabetic patients belong to this group.It is common among individuals of age 45 and above (American Diabetes Association, 2017). Diabetes can cause cardiovascular disease, neuropathy, hypertension, wasting, gangrene, retinopathy, nephropathy, and other functional impairments, all of which raise a person’s chance of death (Liu *et al*., 2016). Other risks include stroke, heart problems, increase in blood pressure, amputation of leg, renal disruption, eyesight loss, and nerve damage. Based on World Health Organization (WHO) statistics, this non-infectious disease (diabetes) has demonstrated salient growth in prevalence with a generational transformation in its global spread in recent years. It is a disease that affects both young and old people in a given population. Diabetes affected close to 422million peopleat the global in the year 2014which reflect a geometric upsurge, as it only affected about 108million individuals in 1980(WHO, 2016). In 2017, 425 million persons (20-79 years old) were predicted to have diabetes, with expectation that the number would have reached 629 million in the year 2045 (Shiferaw & Ayalew, 2018).High expense associated with standard diabetic medicines, as well as their associated side effects, has hampered the chemotherapy and control of diabetes mellitus in modern epoch. Furthermore, most of the diabetic medications tend to have a short-lived effect in the management of diabetes. There is currently a lot of fascination in ethno-medicine that has few or no side effects and of low cost to an average person in society. Currently, plants of medicinal endowment are believed to contain bioactive compoundswhich are efficient in reducing fasting blood glucose baseline with minute toxicity are extensively utilized as alternative therapy in developing nations, despite the fact that the mode of action of this traditional panacea has yet to be fully established scientifically.

## 2. Materials and methodology

### 2.1 choice of chemicals for the work

The set of chemicals and reagents utilized in the experimental design: Hydrochloric acid, hydrogen peroxide, potassium chloride, Tris buffer, sodium hydroxide, sodium carbonate, potassium sodium tartrate,Ellman’s reagent (5,5’-dithiobis-(2-nitrobenzoic acid)), copper sulfate pentahydrate Folin-Ciocalteau reagent, adrenaline, dipotassium hydrogen phosphate trihydrate, potassium dihydrogen phosphate,reduced glutathione, sulfosalicylic acid, trichloroacetic acid, azide sodium chloride,, dipotassium hydrogen orthophosphate. urea, creatinine, glucose test strips, SGOT/AST (Serum Glutamic-Oxaloacetic Transaminase),1-chloro-2,4-dinitrobenzene, alanine aminotransferase (ALT), and lipid profile test kits were obtained from Randox Laboratories, UK. The set of chemical compound used are of high quality and purest analytical grades commercially available.

### 2.2 Plant Materials

Dried calyces of Hibiscus sabdariffa (Zobo leaves), fresh leaves of *Moringa oleifera, Zingiber officinale* (ginger roots), and *Azadirachta indica* (Dogoyaro) leaves were purchased from mile three Market in Port Harcourt River State Nigeria. They were then identified as *Hibiscus sabdariffa* (Zobo), *Moringa oleifera, Zingiber officinale* (ginger roots), and *Azadirachta indica* (Dogoyaro) at the Department of Botany at Rivers State University, Port Harcourt Nigeria.

### 2.3 Animals and experimental design

Forty-two (42) male Wistar rats weighing between 100g to 150g were purchased. The animals were kept in an appropriate plastic enclosure within a well-ventilated animal facility located in Pharmacology Department at Rivers State University, Port Harcourt where they were fed with pellets and water. The animals got housed 10 per plastic cage and exposed to natural period of light of about 12 hours light and 12 hours darkness daily.The animals were categorized into seven groups, each consistingsix animals based on their weights. Prior to treatment, the experimental animals were allowed a week of acclimatization where they were only provided with rat pellets and sufficient water. Subsequent to the week of acclimatization, the animals were then treated for four weeks (28 days) with alloxan and extracts based on their groups.

The arrangement of the experimental groups was outlined below;

**Group 1 (Control):** Animals received their nourished feeds and water orally

**Group 2 (Alloxan):** Animals received 100mg/kg dosage of Alloxanintraperitoneally

**Group 3 (Alloxan + Zobo):** Alloxan-induced diabetic rats were orally administered *Hibiscus sabdariffa* at a dose of 400 mg/kg.

**Group 4 (Alloxan + Moringa):** Alloxan-induced diabetic rats received*M. oleifera*, 200mg/kg through oral route.

**Group 5 (Alloxan + Ginger):** Alloxan-induced diabetic rats were orally given *Zingiber officinale*(500mg/kg).

**Group 6 (Alloxan + Dogoyaro):** Alloxan-induced diabetic rats were orally administered *Azadirachta indica* at a dose of 250mg/kg.

**Group 7 (Alloxan + Glibenclamide):** Alloxan-induced diabetic rats were orally issued Glibenclamide at a dose of 5mg/kg.

### 2.4 Tissue preparation

Cervical dislocation procedure was employed in sacrificing the animals; blood wastaken by puncturing of the eyes using capillary tubes into single sample vials. To obtain serum, the coagulated blood was spin utilizing centrifugal technique at 3000 g for 20 min. The animals’ kidney, hepatocyte tissue (liver) and pancreas were extracted immediately, weighed and processed for biochemical, inflammatory and histological analysis.

### 2.5 Biomakers for liver function test

AST and ALT levels were determined usingLimited accoutrements from Randox Laboratories Reitman and Frankel as shown in the table below

|  | **Solution blank(μl)** | **Specimen(μl)** |
| --- | --- | --- |
| Specimen | ----- | 5 |
| solution 1 | 25 | 25 |
| Distilled water | 5 | ----- |
| Samples were mixed thoroughly and incubation was carried out for 30mins at 37°C |  |  |
| Reagent 2 (R2) | 25 | 25 |
| Samples were Mixed properly and left to settle for a period of 20mins at temperature range of 20 - 25°C |  |  |
| Sodium hydroxide | 250 | 250 |

Samples were mixed and absorbance was takenversus the blank after 5mins at 546nm with a microplate reader and the effect of Aspartate amino transferase(AST) and Alanine amino transferase(ALT) in the serum were extrapolated from a standard curve.

Furthermore, Randox Laboratories Limited accoutrements was used to determine peak position of ALP following the principle characterized by Englehardt as shown in the table below.

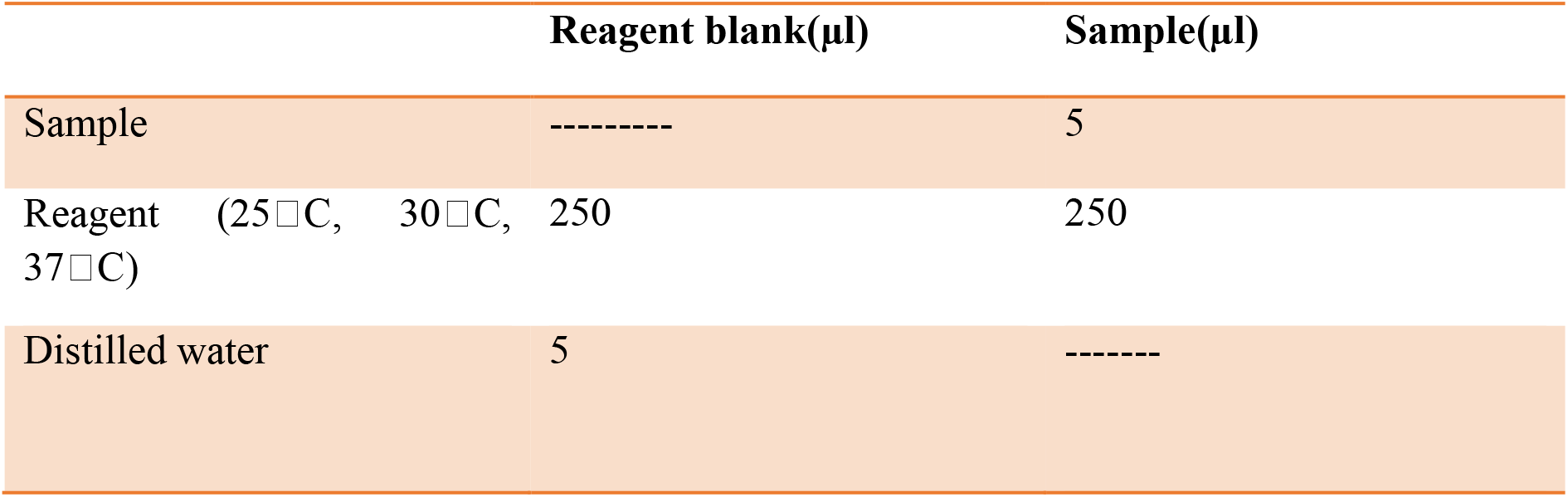

The resultant solutions were mixed and absorbance was read against blank at 405nm with a microplate reader. The timer was then started simultaneously. Readings were taken again exactly after 1, 2 and 3 minutes.

### 2.6 Biomarkers in the kidney function Tests

The concentration of urea was assessed using a commercial kit from randox laboratory. Preparation of the R1 reagent was carried out via combining the contents in R1a and R1b bottles, then by mixing them gently. The composition of R1 and R2 were combined with 660ml and 750ml of water that is distilled, respectively. In each test tube, 10μl of the sample, standard and distilled water were mixed together, and then 100μl of R1 was introducedto each tube. After thorough mixing, incubation was carried out at 37 degree celsius for a period of 10 minutes, thensubsequent adding of 2.5ml of R2 and R3 reagents. The reactant mixtures underwent further incubation at 37°C for 15 minutes. Value of absorbance of the sample A and standard Aversus the blank was measured at 546nm to determine the urea concentration.Concentration of creatinine was also achieved by using randox commercial kit using the procedure shown below

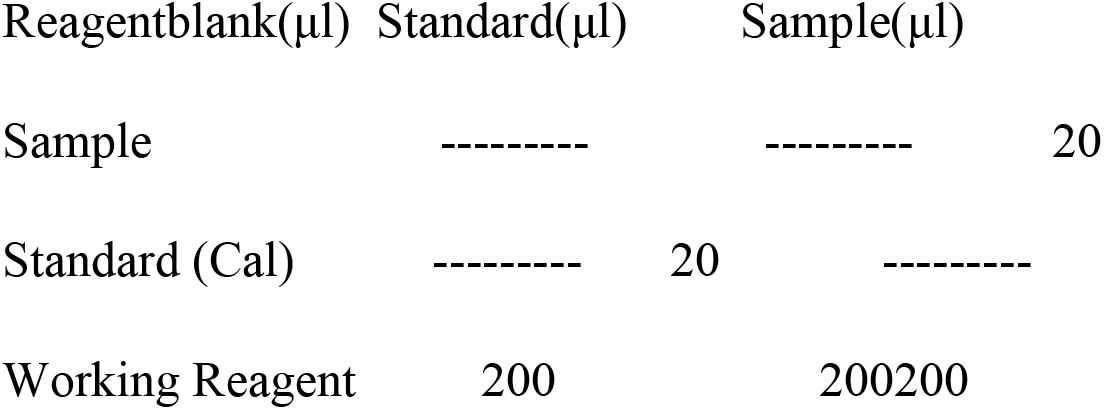
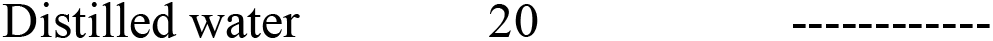

| Reagent blank(μl) | Standard(μl) | Sample(μl) |
| --- | --- | --- |
| Sample | ----- | ----- 20 |
| Standard (Cal) | ----- 20 | ----- |
| Working Reagent | 200 | 200 200 |

The mixing of the reagentswere carried out, and absorbance (A1) of samples and after a period of 30seconds, the result of the standard was determined. The absorbance value of A2 (standard and sample) were determined 2 minutes later at a wavelength of 492nm, then the concentration of creatinine was determined.

### 2.7 Histological examination of liver samples

A portion of the liver from each experimental animal was gutted, flecked and also perfused with 1.15% potassium chloride to remove all traces of haemoglobin which may pollute the apkins. The samples gotten from the liver and the extrahepatic tissues were acquired and preserved in 10% solution of formalin. These apkins were employed for histopathology examination using a routine paraffin-wax embedded system. Sections of 5micrometer consistence was stained with haematoxylin and eosin. The slides got viewed with the aid of microscope for abnormalities.

### 2.8 Statistical analysis

The disparities between the groups were evaluated using the software Graphpad prism. All the generated data were presented as mean value ± standard deviation. Statistical proceedure was performed with one-way analysis of variance (ANOVA). Post hoc testing was conducted using the least significant difference (LSD) and Values of p < 0.05 were of significant consideration.

## 3. Results

### 3.1 Tissue weight

The result of the Table 1 shows a salient decrease(p<0.05) in the relative weight of the body of the alloxan treated organsas compared with the control animals. However, animals treated withherbal teas showed a significant increase (p<0.05) in the relative organ body weight of the animals in all the groupsin comparison with animals in the control group.

**Table 1:** showing the impact of different herbal teas on organ weight of treated rats.

| GROUPS | LIVER | KIDNEY |
| --- | --- | --- |
| GROUP 1 | $2.90 \pm 0.32$ | $1.26 \pm 0.12$ |
| GROUP 2 | $0.22 \pm 0.14$ | $0.74 \pm 0.67$ |
| GROUP 3 | $0.80 \pm 0.35$ | $0.34 \pm 0.41$ |
| GROUP 4 | $0.64 \pm 0.31$ | $0.38 \pm 0.16$ |
| GROUP 5 | $0.72 \pm 0.32$ | $0.34 \pm 0.14$ |
| GROUP 6 | $0.44 \pm 0.20$ | $0.34 \pm 0.14$ |
| GROUP 7 | $0.56 \pm 0.25$ | $0.32 \pm 0.13$ |

Group 1 = Normal Control, Group 2 = Diabetes Control, Group 3 = Zobo plant extract, Group 4 = Moringa extract, Group 5 = Ginger extract, Group 6 = Dogoyaro extract, Group 7 = Glibenclamide.
Values are presented as mean ± standard deviation ( $n=8$ ), $p < 0.05$ against control. Statistical procedure got performed with one-way analysis of variance (ANOVA) and Duncan Post hoc test.

Values are presented as mean ± standard deviation (n=8), p < 0.05 against control. Statistical proceduregot performedwith one-way analysis of variance (ANOVA) and DuncanPost hoc test.

### 3.2 EFFECT OF SPECIFIC HERBAL TEAS ON BLOOD SUGAR LEVELS OF ADMINISTERED ANIMALS

The initial level of glucose was determined in the beginning of the research study for all the rats. Also, the final level of glucose was also obtained at the final stage of the experiment. The starting and final mean glucose levels are shown in figure 1 below. The amount of glucose contained in the the blood of rats induced with diabetes presented a significant upsurge p <0.05 in comparison to the control group rats in all 4 weeks of treatment. While the standard drug attenuated the initial composition of blood glucose of diabetic rats more than the selected herbal teas, the graph shows that over the course of 4 weeks, the herbal teas significantly reduced the blood glucose level of diabetic rats compared to the standard drug. Notwithstanding, compared to rats induced with diabetes alone, administration of herbal teas and standard drug reduced blood glucose levels in the co administered groups. Furthermore, in comparison to rats in the group not treated, administration of selected herbal tea to rats with diabetesdecreased the volume of blood glucose.

**Figure 1.**
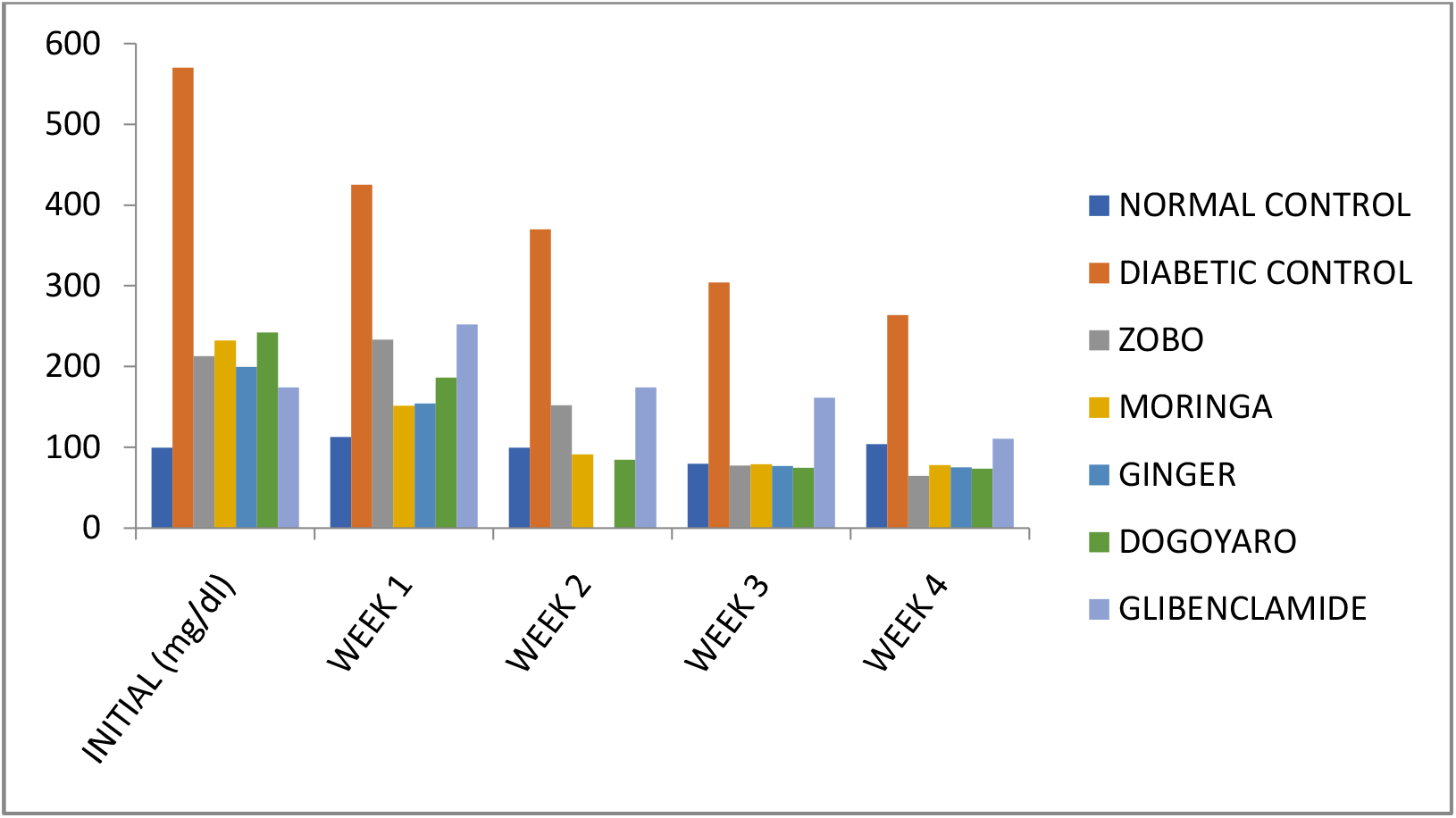
Showing the effectof the various herbal teas on blood sugar(glucose) levels of treated rats.Values are taken as mean ± standard deviation (n=8), p < 0.05 against control. Statistical presentation was carried out using one-way analysis of variance (ANOVA) and Duncan. Post hoc test.

### 3.3 KIDNEY FUNCTION TEST

In comparison to animals in the normal control group, animals treated using all of the selected herbal teas and the standard drug had lower urea and creatinine levels (Figure 2). The results also show that rats induced with diabetes had higher amount of creatinine and by-product of protein metabolism(urea) than rats in the categories of rats without diabetes, and on the other hand, diabetic rats receiving selected herbal teas manifested significant decrease p < 0.05 in urea and level of creatinine, especially for Zobo and Dogoyaro, than rats induced with diabetes alone.

**Figure 2.**
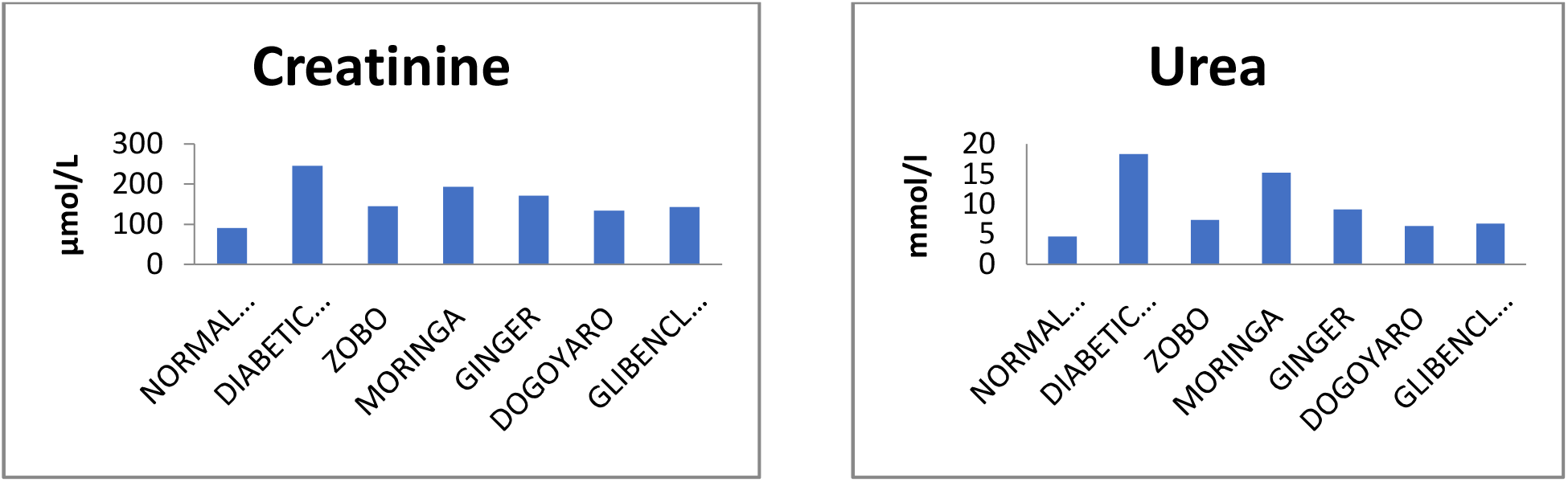
showing the effect of different herbal teas on kidney function biomarkers of treated rats.Results are presented as mean ± standard deviation (n=8), p < 0.05 versus control. One way analysis of variance and Duncan post hoc test were used for the statistical manipulation.

### 3.4 LIVER FUNCTION ASSAY

Rats with induced diabetes had higher expression of AST, ALT, and ALP activities but lower TP and ALB levels in contrast to rats in the normal control rats. The findings additionally demonstrates that diabetic rats co-treated with selected herbal teas exhibited lowerALT, AST, and ALP activities, but showed p < 0.05 of significant elevation in TP and ALBamount than rats induced with diabetes alone. Furthermore, in relation to rats in the untreated group, co-administration of the selected herbal teas in diabetic rats non-significantly reduced the activities of, ALT,AST and ALP while increasing the amount of Total Protein and Albumin in comparison to the rats belonging to the untreated group (figure 3).

**Figure 3.**
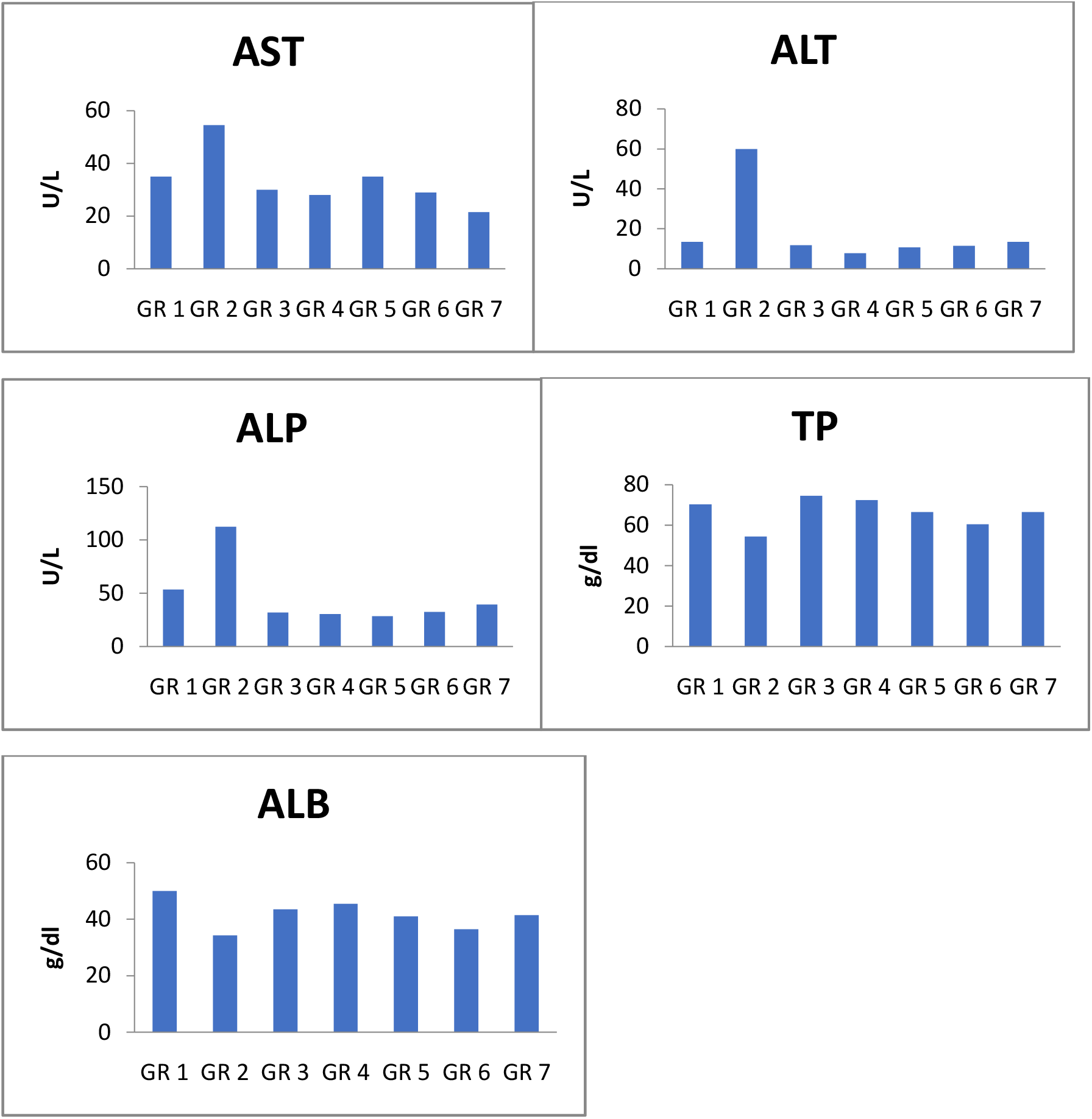
Showing the effect of different herbal teas on liver function test of treated rats.Results are expressed as mean ± standard deviation (n=8), p < 0.05 against control. Statistical analysis was carried out using one-way analysis of variance (ANOVA) and Duncan. Post hoc test.

### 3.5 HISTOLOGICAL ASSESSMENT OF THE LIVER IN TREATED RATS

liver section photomicrographpresents normal Hepatocytes in the control group (figure 4.1) and necrotic hepatocytes in the Alloxan induced diabetic group (figure 4.2). However, co treatment with herbal teas (*Hibiscus Sabdariffa, Zingiber Officinale, Moringa tea, and Dogoyaro*)showed normal Hepatocytes as depicted in figure 4.3-4.7.

Group 1: Normal control

**Figure 4.1.**
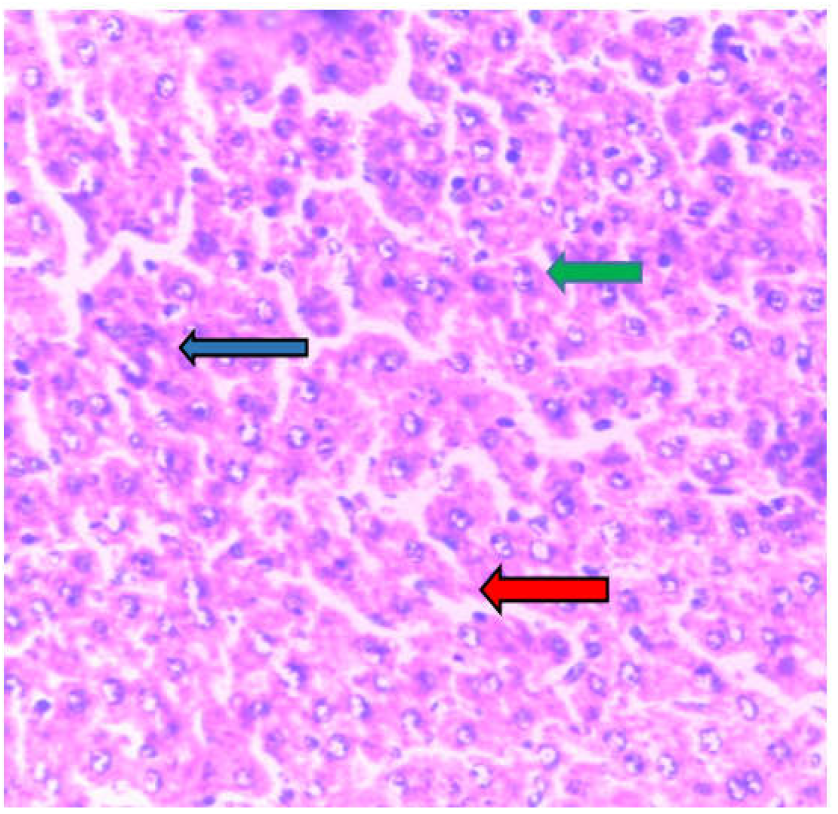
Photomicrograph of the liver section showing normal Hepatocytes (Green), normal portal tract (Blue) and normal central vein (Red) (X100)

Group 2: Diabetic Control

**Figure 4.2.**
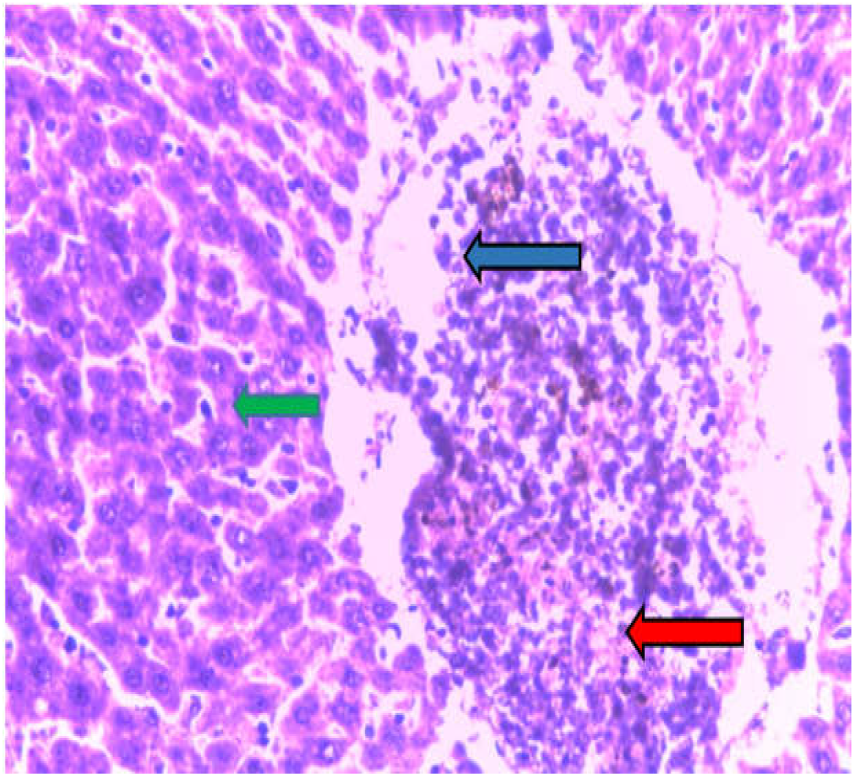
Photomicrograph of the liver section showing necrotic hepatocyes (Red), portal tract infiltrated by lymphocytes (Blue) and congested central vein (Green) (X100)

Group 3: Alloxan + Zobo

**Figure 4.3.**
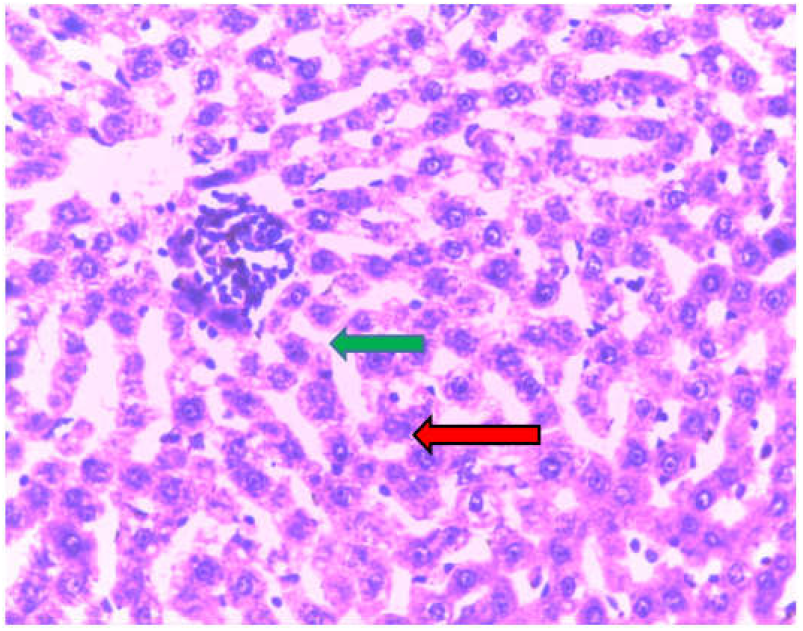
Liver section photomicro-graph showing normal Hepatocytes (Green)and portal tract mildly infiltrated by lymphocytes(Red). (X100)

Group 4: Alloxan + Moringa

**Figure 4.4.**
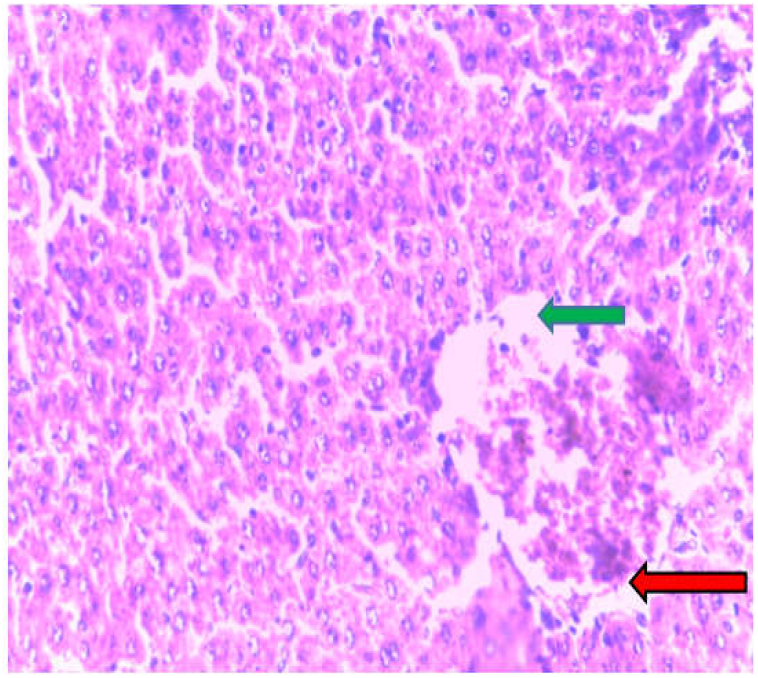
photomicrograph section of the liver showing liver section with normal Hepatocytes (Green) and mildly congested central vein (Red). (X100)

Group 5: Alloxan + Ginger

**Figure 4.5.**
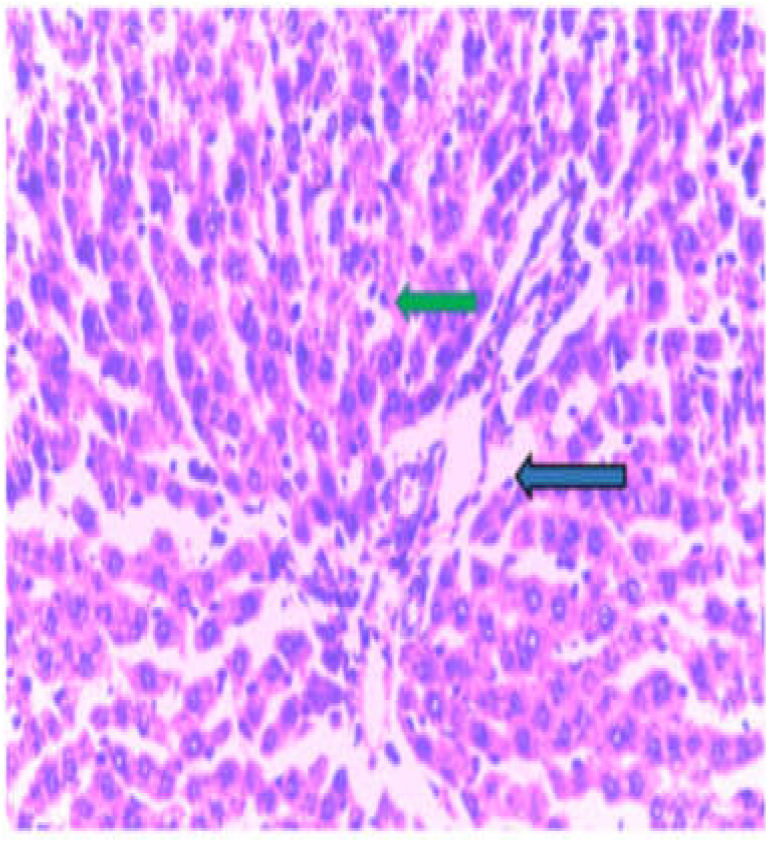
Photomicrograph of the liver section showing normal Hepatocytes (Green)and normal portal tract (Blue).(X100)

Group 6: Alloxan + Dogoyaro

**Figure 4.6.**
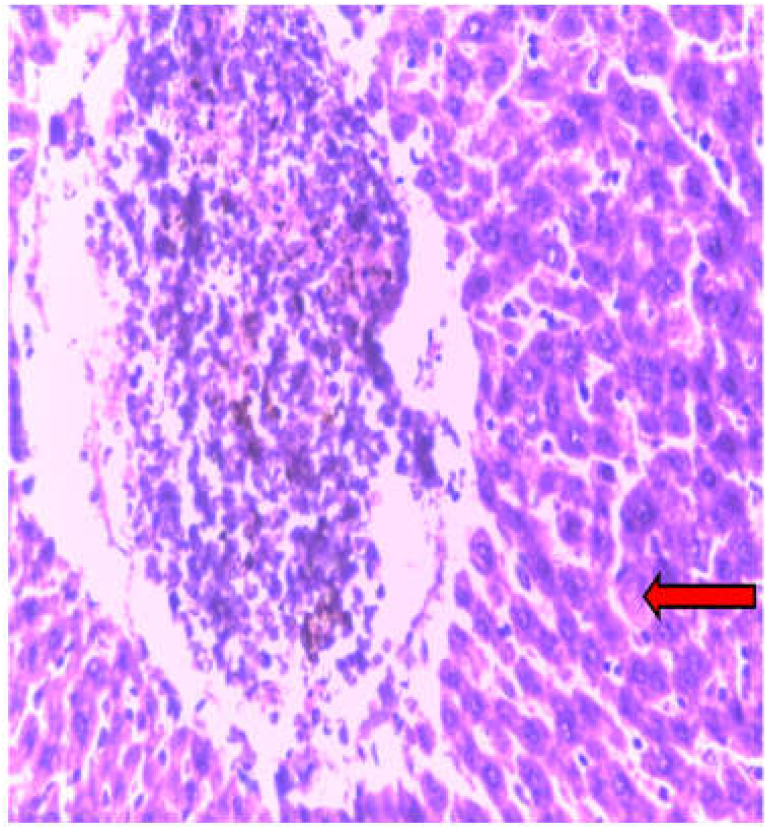
Photomicrograph of the liver section showing mild necrotic hepatocyes (Red). (X100)

Group 7: Alloxan + Glibenclamide

**Figure 4.7.**
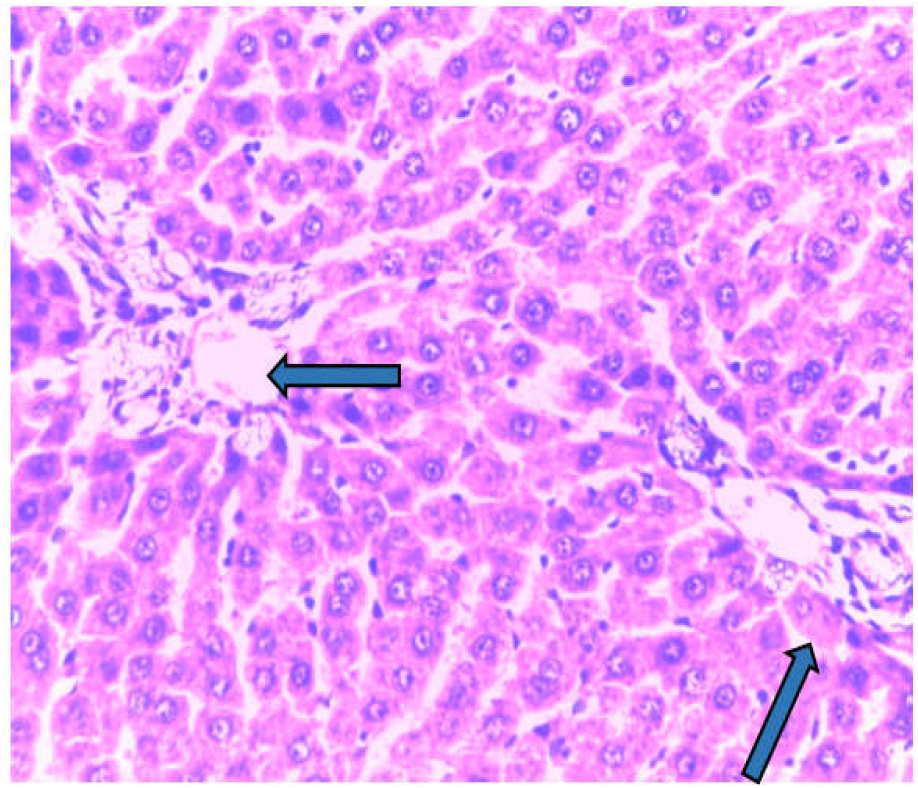
liver section photomicrograph showing normal Hepatocytes (Green)and normal portal tract (Blue). (X100)

## 4.0 DISCUSSION

Diabetes is a persistent hyperglycemic state caused by the failure of the body to make insulin or to use or respond to insulin produced by the body (Ime *et al*., 2022). The study was aimed to see how Zobo tea (*Hibiscus Sabdariffa*), ginger tea (*Zingiber Officinale*), Moringa tea, and Dogoyarotea affected diabetic rats administered alloxan. The current study discovered that alloxan-induced groups had greater blood sugar levels following diabetes induction than untreated control rats, which had normal blood sugar levels during the procedure. However, whereas the concentration of glucose in the systemic circulationof diabetic rats alone remained greatly elevated during the 28-day treatment period, the blood sugarconcentrate indiabetic rats given herbal teas, glibenclamide decreased demonstrating extracts’ potential in controlling hyperglycemia in diabetes. The output of this research work supports the finding of thepast studies (Ayobami *et al*., 2020; Ime *et al*., 2022). Also, though the difference in the amount of blood glucose in animal(rats) administered with teas extracted from the plants was not significant, it was discovered that Zobo tea performed slightly better than the others, followed by Dogoyaro tea, Moringa tea, and Ginger tea.

Alloxan induced diabetes also brought about a significant loss in liver and kidney weight in animals in comparison to the normal control (Table 1). It was discovered that among all teas extracted from plants, *Azadirachta indica* (Dogoyaro tea) improved organ weight loss the least, as did the standard drug. While *Hibiscus Sabdariffa* (Zobo tea) mitigated liver weight loss more compared to the other teas (*Zingiber Officinale, Moringa*, and*Azadirachta indica*), Moringa tea mitigated kidney weight loss more compared to the other teas.

Elevated serum levels of liver markers indicate liver damage. The rats with diabetes induced by alloxan in this study had elevated AST, ALT and ALP activities compared with normal control mice, demonstrating diabetes-induced liver damage.This finding is in tandem with the report of Elizabeth(2005) who reported a higher incidence of liver function abnormalities among people with type 2 diabetes than individuals without diabetes. Total albumin and protein levels of diabetic rats injected with alloxan were reduced in comparison with normal control mice that is in agreement with the result ofOmodanisi*et al*., 2017 with a report of a great reduction in total protein, albumin and globulin concentration in diabetic rats compared to non-diabetic rats. However, administration of *Hibiscus Sabdariffa, Zingiber Officinale, Moringa* and*Azadirachta indica* tea extracts to rats with diabetes for a period of four weeks timeframe resulted in a substantial decrease in serum AST, ALT, ALP functions and an increase in total protein and albumin compared with diabetic control rats which also agrees with the finding of Omodanisi*et al*., 2017 which suggested that Moringa Oleifera-based feed supplement could hinder I/R-induced kidney injury and reduce oxidative stress. This effect suggests the hepatoprotective potential of Hibiscus Sabdariffa, Zingiber Officinale, Moringa and Azadirachta indica. Furthermore, our histological assessment showed the therapeutic potentials of these selected herbal teas owning to their ability to revert necrotic hepatocytes induced by Alloxan.

The present study also showed diabetic kidney damage, as evidenced by elevated amount of urea and creatinine in rat induced with alloxan having diabetesin comparison with those not treated,which is in tandem with a 2008research reported in the Journal of Nepal Association for Medical Laboratory Sciences, which affirms that people with diabetes were deduced to show significantly higher amount of creatinine and urea than healthy individuals. Diabetic rats given selected herbal tea extracts showed substantially decreased levels of kidney function (urea level, creatinine level) compared to rats with diabetes alone.

## 5.0 CONCLUSION

The study indicated a potential protective role of Zobo tea (*Hibiscus Sabdariffa*), ginger tea (*Zingiber Officinale*), Moringa tea, and Dogoyaro tea inmaintaining blood sugar levels and abating nephro-hepatotoxicity, as evident by differences in the amount of sugar contained in the blood,kidney and liver function markers alongside withPhotomicrograph of the liver sections.

## Acknowledgement

Prof A.S. Ezekwe of Rivers State University Nigeira, for his guide all through this work.

## Contribution of Authors

EN., ISM., added to the conceptualization and modeling of the work. ES., ISM., VDS., EAO. and MOI. were all involved in the laboratory experimentation EN., ISM., VDV., were involved in the interpretation of data. EN., ISM., EAO and MOI drafted the manuscripts.

## Ethical Approval

All animals used in this study, were handled strictly in accordance with the River State University Nigeria Ethics Committee guidelines for use of experimental animal research

